# Neuronal properties of pyramidal cells in lateral prefrontal cortex of the aging rhesus monkey brain are associated with performance deficits on spatial working memory but not executive function

**DOI:** 10.1101/2023.02.07.527321

**Authors:** Tara L. Moore, Maria Medalla, Sara Ibañez, Klaus Wimmer, Chromewell A. Mojica, Ronald J. Killiany, Mark B. Moss, Jennifer I. Luebke, Douglas L. Rosene

**Affiliations:** Department of Anatomy & Neurobiology, Boston University Chobanian & Avedisian School of Medicine, Boston, MA USA 02118; Center for Systems Neuroscience, Boston University, Boston, MA USA 02118; Centre de Recerca Matemàtica, Edifici C, Campus Bellaterra, 08193 Bellaterra, Spain

## Abstract

Age-related declines in cognitive abilities occur as early as middle-age in humans and rhesus monkeys. Specifically, performance by aged individuals on tasks of executive function (EF) and working memory (WM) is characterized by greater frequency of errors, shorter memory spans, increased frequency of perseverative responses, impaired use of feedback and reduced speed of processing. However, how aging precisely differentially impacts specific aspects of these cognitive functions and the distinct brain areas mediating cognition are not well understood. The prefrontal cortex (PFC) is known to mediate EF and WM and is an area that shows a vulnerability to age-related alterations in neuronal morphology. In the current study, we show that performance on EF and WM tasks exhibited significant changes with age and these impairments correlate with changes in biophysical properties of L3 pyramidal neurons in lateral LPFC (LPFC). Specifically, there was a significant age-related increase in excitability of Layer 3 LPFC pyramidal neurons, consistent with previous studies. Further, this age-related hyperexcitability of LPFC neurons was significantly correlated with age-related decline on a task of WM, but not an EF task. The current study characterizes age-related performance on tasks of WM and EF and provides insight into the neural substrates that may underlie changes in both WM and EF with age.

## Introduction

The cognitive domain of Executive Function (EF) consists of abilities such as abstraction, cognitive flexibility, planning, shifting of response set and inhibition of perseveration.^1, 2^ Working Memory (WM), a component of the cognitive domain of memory is highly related to EF and was first described by Baddeley and Hitch in 1974^3^, It consists of the ability to retain and operate on information over a short period of time.^4-7^ Both EF and WM are critical for decision making, information processing, learning and adaption, all of which are necessary for activities of daily life.^1, 2, 4^ Age-related declines in EF and WM occur in humans and rhesus monkeys^8-17^ as early as middle-age. This is evident by declines in performance on classic tests of EF and WM including the Wisconsin Card Sorting Task, the Stroop Task, Delayed Response, Reversal Learning, Delayed Recognition Span Task and the Category Set Shifting Task where the severity of impairment increases with advancing age.^15, 17-22^ Specifically, performance by aged individuals on these tasks is characterized by greater frequency of errors, shorter memory spans, increased frequency of perseverative responses and impaired use of feedback and speed of processing.

The prefrontal cortex (PFC), specifically its lateral subdivision, is thought to mediate EF and WM and is known to change with age. While there is no significant loss of neurons in the PFC,^23, 24^ studies in rhesus monkeys have demonstrated age-related decreases in grey and white matter volume, degenerative changes in myelin and decreased level of monoamines and their receptors specifically within lateral PFC (LPFC) including area 46 ^19, 23, 25-34^. In addition, a loss of myelin integrity in area 46 and underlying frontal white matter, as measured by electron miscopy and by decreased fractional anisotropy using *in vivo* diffusion MR imaging in monkeys, has been shown^29^ There is an overall decrease in the volume of white matter with age, which is most prominent in the frontal lobe.^35-38^ Taken together, these findings show the vulnerability of LPFC to age-related alterations in morphology that are associated with declines in cognitive function. However, despite the well-established changes in EF and WM with age, the temporal progression of precise changes in the PFC that are associated with distinct aspects of EF and WM deficits remain largely unknown.

Work from our group and others have shown age-related changes in neuronal properties of the LPFC.^36-51^ Specifically, *in vitro* electrophysiological studies of single-neuron biophysical properties have demonstrated that layer 3 (L3) pyramidal neurons in LPFC exhibit hyperexcitability with age, associated with increased input resistance and increased action potential firing frequency in response to step current injections. ^39, 40, 52^ Further, with both the electron microscopy and electrophysiology, we and others have demonstrated significant age-related decline in excitatory spines, synapses and synaptic currents in LPFC neurons.^23, 36, 38, 41-45, 53-56^ Importantly, these sub-lethal age-related changes in L3 LPFC neurons have been shown to strongly correlate with age-related cognitive decline, especially with impairments in the Delayed Recognition Span Task – Spatial (DRSTsp, Fig. 1A,B); a task of spatial working memory task.^42, 57^ In a recent computational modeling study of the DRSTsp task, we have shown that age-related increases in AP firing of L3 pyramidal neurons in LPFC are associated with age-related impaired maintenance of sequence information, and in turn performance, on the DRSTsp task.^57^ Indeed, a full dynamic range of AP firing frequencies together with facilitation of excitatory synaptic transmission, likely from recurrent collaterals, of LPFC L3 pyramidal neurons, are predicted to be the most important neuronal properties that support DRSTsp performance.^57^ However, it remains unclear how LPFC neuronal properties relate to other aspects of EF and specifically to abilities such as abstraction, set-shifting and perseveration, which are critical for performance on our EF task, the Category Set Shifting Task (CSST, Fig. 1C).

**Figure 1.**
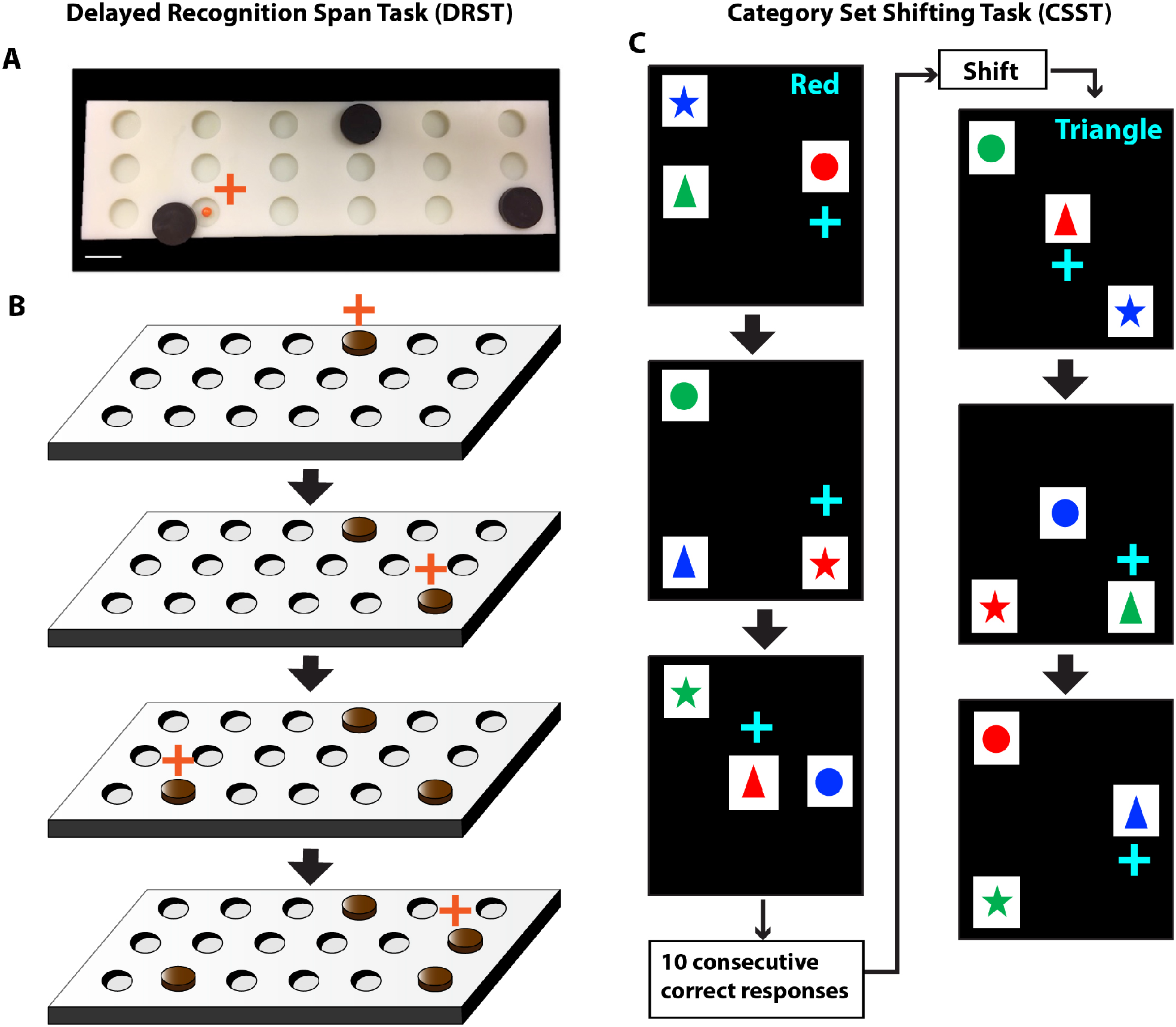
Schematic of Delayed Recognition Span Task and Category Set Shifting Task. A) A photograph of the testing board and the sample discs used in the DRST. B) A schematic showing sequential stimuli (brown disc) presentation within one trial of the DRST. During each presentation the monkey must choose the disc in the new spatial location. Each successive correct response trial was followed by the addition of a new disc in a novel location on the testing board and this continued until the monkey made an error (i.e., chose a previously chosen disc).With the occurrence of the first error, the trial was terminated and the number of discs on the testing board minus one were counted to determine the recognition span score for that trial (i.e., number of correct consecutive responses). C) In this schematic of the Category Set Shifting Task (CSST) each black screen (panel) represents one trial. On each trial of the CSST, the monkey is presented with three stimuli that vary in shape and color. During the first concept condition, the monkey must choose the red stimulus regardless of its shape as illustrated sequentially in the top three screens of this figure. Once the monkey chooses the correct stimulus on 10 consecutive trials, the computer switches the rewarded stimulus on the same testing day, without alerting the monkey. In the second concept condition, the monkey must choose the triangle shaped stimulus, regardless of the color as illustrated in the bottom three screens of the figure. Again, when the monkey chooses the correct stimulus for 10 consecutive trials the computer switches the rewarded stimulus on the same testing day, without alerting the monkey. Testing is continued in this same manner for the blue and star concept conditions.

In the current study, we compared performance by young, middle-aged and aged rhesus monkeys on both the Delayed Recognition Span Task - Spatial (DRSTsp, working memory) and the Category Set Shifting Task (CSST, executive function). We show how performance on these two tasks correlates with biophysical properties of LPFC L3 pyramidal neurons. While the DRSTsp task used in our previous study^57^ provides a measure of spatial working memory, the CSST, modelled after the human Wisconsin Card Sorting Test, assesses EF features including abstraction, set shifting, response maintenance and preservation, and therefore provides an broader assessment of EF abilities. ^8, 9, 58^ The current study characterizes age-related performance on tasks of WM and EF and provides insight into the specific neural substrates that may underlie changes in both WM and EF with age.

## Methods

### Subjects

Behavioral data for this study was collected from 74 rhesus monkeys (Macaca mulatta) of both sexes between the ages of 5-30 years using identical testing protocols. Based on an extensive survival study at Emory Regional Primate Research Center, which suggests a ratio of three to one between monkey and human years of age, we have classified young monkeys as between 5 and12 years of age, middle-aged monkeys as between 13 and 20 years of age and aged monkeys as > 20 years old^59^. As detailed in Table 1, data was collected from 16 young monkeys (nine males and seven females), 33 middle-aged monkeys (15 males and 18 female) and 25 aged monkeys (13 males and 12 females) as detailed in Table 1. All monkeys had known birth dates, complete health records and were obtained from National Primate Research Centers or private vendors. All monkeys received medical examinations before entering the study. In addition, explicit criteria were used to exclude monkeys with a history of any of the following: splenectomy, thymectomy, exposure to radiation, cancer, organ transplantation, malnutrition, chronic illness including viral or parasitic infections, neurological diseases or chronic drug administration. Each of the monkeys underwent magnetic resonance imaging (MRI) to ensure there was no overt neurological damage. Results of the evaluations revealed that all monkeys were healthy at the time they were entered into the study.

**Table 1.** List of Subjects and Sex, Age Group and Age at Testing.

| ID | Sex | Age Group | Age at CSST | Age at DRST | Monkeys with Electrophysiological Data |
| --- | --- | --- | --- | --- | --- |
| AM202 | F | Y | 10.1 | 9.4 | N |
| AM254 | F | Y | 10.7 | 9.1 | Y |
| AM095 | F | Y | 7 | 6.2 | N |
| AM163 | F | Y | 9.7 | 9.1 | N |
| AM195 | F | Y | 11.8 | 10.9 | N |
| AM214 | F | Y | 10.1 | 9.3 | N |
| AM255 | F | Y | 10 | 7.3 | Y |
| AM204 | M | Y | 5.9 | 6.3 | N |
| AM230HL | M | Y | 6.8 | 6.6 | N |
| AM229HL | M | Y | 7.1 | 6.9 | N |
| AM222 | M | Y | 7.2 | 6.3 | N |
| AM132 | M | Y | 7.2 | 6.4 | N |
| AM367 | M | Y | 8.2 | 7.6 | Y |
| AM296 | M | Y | 8.4 | 7.9 | Y |
| AM352 | M | Y | 9.4 | 9.2 | Y |
| AM295 | M | Y | 10.4 | 9.9 | Y |
| AM291 | F | MA | 13.3 | 13 | N |
| AM278 | F | MA | 14.8 | 13.7 | Y |
| AM251 | F | MA | 16.7 | 16.5 | N |
| AM250 | F | MA | 16.9 | 16.6 | N |
| AM257 | F | MA | 17 | 15.2 | N |
| AM297 | F | MA | 17.4 | 17 | Y |
| AM253 | F | MA | 17.9 | n/a | N |
| AM161 | F | MA | 18.7 | 18.2 | N |
| AM252 | F | MA | 18.8 | 17.2 | N |
| AM293 | F | MA | 19.2 | 18.7 | N |
| AM256 | F | MA | 19.9 | 18.4 | N |
| AM212 | F | MA | 20 | 19.3 | N |
| AM160 | F | MA | 20.5 | 19.6 | N |
| AM287 | F | MA | 20.6 | 20.2 | N |
| AM126 | F | MA | 20.4 | 20.1 | N |
| AM149 | F | MA | 19.2 | 18.4 | N |
| AM159 | F | MA | 19.4 | 18.6 | N |
| AM190 | F | MA | 17.8 | 17.2 | N |
| AM353 | M | MA | 14.3 | 14 | N |
| AM034L | M | MA | 15.2 | n/a | N |
| AM143 | M | MA | 15.5 | 14.7 | N |
| AM288 | M | MA | 15.6 | 15.1 | Y |
| AM158h | M | MA | 16.9 | 15.7 | N |
| AM223L | M | MA | 17.5 | 16.8 | N |
| AM274 | M | MA | 17.7 | 16.6 | Y |
| AM233H | M | MA | 17.9 | 17.7 | N |
| AM209L | M | MA | 18.2 | 18.1 | N |
| AM279 | M | MA | 19 | 18.5 | Y |
| AM124 | M | MA | 19.2 | 19 | N |
| AM133 | M | MA | 19.2 | 19 | N |
| AM226L | M | MA | 19.9 | 19 | N |
| AM314 | M | MA | 20.2 | n/a | Y |
| AM123 | M | MA | 20.4 | 20.2 | N |
| AM179 | F | Aged | 23.2 | 22.8 | N |
| AM162 | F | Aged | 22 | 21 | N |
| AM286 | F | Aged | 21.5 | 20.7 | Y |
| AM196 | F | Aged | 22.3 | 21.2 | N |
| AM234 | F | Aged | 23 | 22.2 | N |
| AM235 | F | Aged | 24.2 | 23.6 | N |
| AM090 | F | Aged | 24.4 | 23.6 | N |
| AM236 | M | Aged | 21.5 | 20.9 | N |
| AM208 | M | Aged | 22.5 | 21.4 | N |
| AM281 | M | Aged | 22.5 | 22.2 | Y |
| AM282 | M | Aged | 22.6 | 22.4 | Y |
| AM276 | M | Aged | 23.6 | 22.7 | N |
| AM243 | M | Aged | 24 | 22.2 | N |
| AM024L | F | Aged | 29.4 | n/a | N |
| AM371 | F | Aged | 25.8 | 25.5 | Y |
| AM275 | F | Aged | 27.1 | 26.3 | N |
| AM370 | F | Aged | 27.2 | 26.6 | N |
| AM098 | F | Aged | 27.7 | 26.9 | N |
| AM284 | M | Aged | 25.1 | 24.3 | Y |
| AM283 | M | Aged | 26.1 | 25.3 | Y |
| AM304 | M | Aged | 26.1 | 25.9 | N |
| AM109 | M | Aged | 26.4 | 25.7 | N |
| AM298 | M | Aged | 26.5 | 25.9 | Y |
| AM091 | M | Aged | 30 | 29.7 | N |
| AM374 | M | Aged | 26 | 25.3 | N |

Rows highlighted in gray indicate monkeys with both cognitive and electrophysiological data. Brains were harvested between 2-7 months following the completion of cognitive testing.

While on study, monkeys were individually housed in colony rooms in the Boston University Animal Science Center where they were in constant auditory and visual range of other monkeys. This facility is fully AAALAC accredited, and animal maintenance and research were conducted in accordance with the guidelines of the National Institutes of Health and the Institute of Laboratory Animal Resources Guide for the Care and Use of Laboratory Animals. All procedures were approved by the Boston University Institutional Animal Care and Use Committee. Diet consisted of Lab Diet Monkey Chow (#5038 - LabDiet Inc., St. Louis, MO) supplemented by fruit and vegetables with feeding taking place once per day, immediately following behavioral testing. All monkeys were fed 12– 20 biscuits per day based on their weight. During testing, small pieces of fruit or candy were used as rewards. Water was available continuously. The monkeys were housed under a 12-h light/dark cycle with cycle changes occurring in a graded fashion over the course of an hour. They were checked daily by trained observers for health and well-being and were given a medical exam every three months by a Clinical Veterinarian in the Boston University Animal Science Center.

### Cognitive Testing

The monkeys in this study were part of a larger study of normal aging and were behaviorally sophisticated, having experience with the Delayed Non-Matching to Sample task prior to the administration of the Delayed Recognition Span Task and the CSST.^10, 14, 17, 18, 58^ For the current study, we are presenting the cognitive data from the DRSTsp, a task of spatial working memory and from the CSST, a task of abstraction and set shifting. For all tasks, white noise was played on two speakers located within the automated apparatus to mask extraneous sounds. A non-correctional procedure was used with small pieces of candy as rewards.

### Delayed Recognition Span Task

The DRST was administered in a Wisconsin General Testing Apparatus (WGTA) and the testing board had three rows of six wells each (3.5 cm wide, 0.5 cm deep) and with the wells spaced 6 cm mm apart within a row. (Fig.1A,B) and the rows were spaced 1.5 cm apart. For this task, 15 identical plain brown discs (6 cm in diameter) were used as stimuli. During the first sequence of a trial, 1 disc was placed over 1 of the 18 wells, which was baited with a food reward. The WGTA door was raised and the monkey was allowed to displace the disc to obtain the reward. The door was then lowered, the first disc was returned to its original position over the now unbaited well, and a second disc was placed on the board over a baited well in a different spatial location. After 10 s, the door was once again raised, and the monkey was required to identify the new second disc in its novel spatial location to obtain the reward. Each successive correct response trial was followed by the addition of a new disc in a novel spatial location on the testing board and this continued until the monkey made an error (i.e., chose a previously chosen disc). With the occurrence of the first error, the trial was terminated and the number of discs on the testing board minus one were counted to determine the recognition span score for that trial (i.e., number of correct consecutive responses). Ten such trials were presented each day for 10 days (total 100 trials).

### Category Set Shifting Task

An automated pretraining task (the three-choice discrimination task) followed by the CSST were sequentially administered in an automated General Testing Apparatus containing a touch sensitive, resistive, computer screen, driven by a Macintosh computer (1.83 GHz Intel Core 2 Duo Processor). The testing apparatus had a darkened interior and was in a darkened room.

An automated pre-training task was used to teach the monkey to touch the computer screen.^58^ The pre-training task required the monkey to touch a single stimulus which appeared in one of 9 random locations on the screen to receive a food reward. This was continued for 20 trials a day until the monkey correctly responded for 20 consecutive trials in a single day. If a monkey did not respond within one minute, the screen reverted to black, a non-response was recorded, and the intertrial interval began followed by the next trial. The intertrial interval for each trial was 15 seconds. The day after the monkey completed the pre-training task, monkeys began a simple three-choice discrimination task. This task was administered to determine if there was a difference in the performance across the age groups in discriminating among three fixed stimuli based on the reward contingency. The task presented the monkey with a pink square, orange cross and a brown 12 point star. The stimuli remained constant in terms of color and shape for each trial but appeared in a pseudo-random order in 9 different spatial locations on the screen for 80 trials per day. The pink square was the positive stimulus for all trials and all monkeys. A non-correctional procedure was used and the monkey was given a food reward only when he/she correctly touched the pink square on the screen. To reach criterion, the monkey had to choose the pink square on ten consecutive trials during one testing session.

Next, formal testing began on the CSST. Each day of testing consisted of 80 trials where three stimuli appeared in three of nine pseudo-random locations on the computer touch screen, as shown in Fig. 1C. The stimuli differed in two relevant dimensions, color (red, green, or blue) and shape (triangle, star, and circle). Each color and each shape was presented on every trial with all nine possible combinations of stimuli (i.e. red triangle, red star, red circle, blue triangle, etc.) presented in a pseudo-random sequence in a balanced fashion over four days of testing. If a monkey did not respond within one minute, the screen reverted to black, a non-response was recorded, and the intertrial interval began followed by the next trial. The intertrial interval for each trial was 15 seconds.

For the first abstraction, red was designated as the positive dimension and the monkey had to choose the red stimulus regardless of its shape to obtain a food reward. Once the monkey chose this stimulus on ten consecutive trials the program switched the rewarded contingency during the same testing session, without alerting the monkey. Now, the monkey had to choose the stimulus shaped like a triangle, regardless of its color, to obtain a food reward. Again, when the monkey reached a criterion of 10 consecutive responses, the computer switched the rewarded contingency so that the blue stimulus then had to be chosen, regardless of its shape, to obtain a food reward. Finally, when criterion was reached on the blue category, the contingency was switched to the last category, star.

### Perfusion and Tissue Biopsy

Monkeys were euthanized approximately 2-7 months after completing cognitive testing. Brains were perfused using our two-stage Krebs-Paraformaldehyde perfusion method for harvesting live tissue and subsequent fixation^52, 60-62^. The monkeys were initially sedated with Ketamine hydrochloride (10 mg/ml, IM) and deeply anesthetized with sodium pentobarbital (to effect, 15 mg/kg, IV), and a craniotomy performed over the left hemisphere. Then the chest was opened, the ascending aorota cannulated and the brain perfused beginning with ice-cold Krebs-Henseleit buffer (mM: 6.4 Na2HPO4, 1.4 Na2PO4, 137 NaCl, 2.7 KCl, 5 Glucose, 0.3 CaCl2, 1 MgCl2; pH 7.4, 4°C). Within ten minutes of opening of the chest cavity (anoxia), a block of tissue (1 cm^3^) that included the ventral bank of LPFC was removed and transferred to oxygenated (95% O2, 5% CO2) ice-cold Ringer’s solution (mM: 26 NaHCO3, 124 NaCl, 2 KCl, 3 KH2PO4, 10 glucose, 1.3 MgCl2, pH 7.4), and sectioned into 300-μm coronal slices with a vibrating microtome. Slices were collected in oxygenated room temperature Ringer’s solution. Once fresh tissue harvesting was concluded, perfusate was switched to freshly depolymerized 4% paraformaldehyde in 0.1 M phosphate buffer (PB, pH 7.4, at 37°C) to fix the intact hemisphere and the remaining brain tissue.

### Whole Cell Patch-Clamp Recording and Assessment of Electrophysiological Properties

After 1hr equilibration in oxygenated Ringer’s solution, acute slices were placed into submersion-type recording chambers (Harvard Apparatus, Holliston, MA, USA) and visualized under Nikon E600 infrared-differential interference contrast (IR-DIC) microscopes. Standard tight-seal, whole-cell patch-clamp recordings with simultaneous biocytin filling were obtained from layer 3 pyramidal cells of LPFC, as previously described ^39, 52, 63, 64^. Patch electrodes were fabricated on a horizontal Flaming and Brown micropipette puller (Model P-87, Sutter Instruments, Novato, CA, USA). Potassium methanesulfonate-based internal solution (mM: 122 KCH3SO3, 2 MgCl2, 5 EGTA, 10 NaHEPES, with 1% biocytin, pH 7.4), with resistances of 3-6 MΩ in the external Ringer’s solution was used for recording. Data were acquired using EPC-9 or EPC-10 patch-clamp amplifiers using PatchMaster software (HEKA Elektronik, Lambrecht, Germany). Bessel filter frequency was 10 kHz and sampling frequency was at 7 kHz for voltage clamp and 12 kHz for current clamp recordings. The series resistance ranged from 10-15 MΩ and was not compensated. All physiological experiments were conducted at room temperature, in oxygenated Ringer’s solution (superfused at 2-2.5 ml/min), which improves the viability and duration of recordings from monkey slices.

Neurons verified to reside in L3 were selected for electrophysiological analyses based on well-established inclusion criteria: A resting membrane potential ≤ -55 mV, stable access resistance, action potential (AP) overshoot and repetitive firing responses (Amatrudo et al., 2012). Passive membrane properties, resting membrane potential (Vr), input resistance (Rn), and membrane time constant (tau) were measured using a series of 200 ms depolarizing and hyperpolarizing current steps ^39, 52, 63, 64^. Input resistance (Rn) was calculated as the slope of the best-fit line of the voltage-current linear relationship. Membrane time constant (tau) was measured by fitting a single exponential function to the membrane potential response to a -10 pA hyperpolarizing current step. Single AP firing properties (threshold, amplitude, rise time, fall time, and duration at half maximal amplitude) were measured from the second AP in a train of 3 or more spikes generated by the smallest current step. Rheobase was measured as the minimum current required to evoke a single AP during a 10 sec depolarizing current ramp stimulus (0–200 pA). A series of 2 s hyperpolarizing and depolarizing current steps (−170 to +380 pA, using either 20 or 50 pA increments) was used to assess active and repetitive firing properties. Traces were exported to Matlab to assess electrophysiological properties.

### Outcome Measures

#### DRST

The mean total span achieved by each monkey was determined across the 10 trials per day for 10 days of testing.

#### CSST

The total number of trials and errors to criterion for the 3 choice discrimination task, the total number of trials and errors to criterion for the red condition and the total number of trials, errors and perseverative errors for the three shift conditions (triangle, blue and star) were determined. Because the aging monkeys overall required more trials to reach criterion, it is plausible that the analysis of total perseverative errors was skewed by the increased number of opportunities that the aging monkeys had to make perseverative errors. Therefore, to control for the different number of trials we further analyzed this type of error by calculating the total perseverative errors as a percentage of total shift trials. In addition, the total number of broken sets and the total number of non-responses were determined. A perseverative error was recorded when a monkey made an error by choosing a stimulus that contained a component of the previously rewarded concept. A broken set was recorded when a monkey achieved a span of six to nine consecutive correct responses and then made an error, just missing criterion. A non-response was recorded when a monkey failed to respond by touching the screen on any trial within one minute of the stimuli appearing on the screen. A non-response was also recorded as an error and therefore the total number of consecutive correct responses was reset to zero when a non-response occurred.

#### Electrophysiology

The outcome measures that estimate passive and active firing properties were obtained from each individual neuron and compared across age groups in a repeated measures design. Passive membrane properties include: resting membrane potential (Vr), input resistance (Rn), membrane time constant (tau). Active properties include: Rheobase (minimum current to elicit an AP); repetitive AP firing frequency in response to 2-second depolarizing current steps (−170 to +380 pA, using 50 pA increments).

#### Data Analysis

All analyses were performed in Matlab R2022a. We first assessed the data for a sex effect on any of the outcome measures. This analysis revealed no effect of sex on any measure so data for males and females were pooled for all analyzes. We used standard linear regressions (“fitlm” function) to determine the correlation between age and performance on the DRST and between age and performance on the CSST (treating age as a continuous variable), the correlation between the firing rate and performance on the DRST and on the CSST, and the correlation between the input resistance and the firing rates. In the analyses of the DRST and CSST performance variables (one data value per subject) vs. firing rate, the firing rates were first averaged within each subject.

Then, to further analyze the effect of age, treated as a categorical variable (young, middle-aged and aged groups), on performance on the DRST and CSST, we performed separate one-way ANOVAs (“anova1” function in Matlab) for total span on the DRST, and for CSST total trials and errors on the initial abstraction, total perseverative errors, perseverative errors as a percent of shift trials, broken sets and non-responses. In addition, a two-way repeated measures ANOVA, including age group and task condition, was also performed for the trials and errors on the three shift conditions (“fitrm” and “ranova” functions). A two-way repeated measures ANOVA was conducted to determine whether there were differences in firing rate vs. age group, with injected current as the repeated measure. After the ANOVAs, *post-hoc* analyses were conducted using Tukey’s honestly significant difference procedure of estimated marginal means for multiple comparisons (“multcompare” function).

Linear mixed-effects models with random effects were used to determine the relationship between age and the biophysical intrinsic and firing properties of the neurons (“fitlme” function). Subjects were treated as random effect blocking factors,^65, 66^ because electrophysiological data were collected from several neurons per subject.

### Mediation analysis

A mediation analysis was conducted to examine whether the effect of age on DRST task performance was mediated by a change in prefrontal AP firing rate. Mediation analysis assesses whether covariance between predictor and dependent variable is explained by a third mediator variable. Significant mediation is obtained when inclusion of the mediator (indirect effect) in the model significantly alters the slope of the predictor-dependent variable relationship (direct effect). To carry out the analysis, we used the Multilevel Mediation and Moderation (M3) Toolbox and tested the significance using bootstrap, producing two-tailed p-values.^67^

For all analyses, the significance level was α = 0.05.

## Results

### Delayed Recognition Span Task Performance

With age as a continuous variable, we found that the total spatial span on the DRST significantly decreased with age (Fig. 2A; Table 2, F(1,69) = 14.3, **p = 3.4 × 10**^**-4**^). These data are consistent with our previous finding that DRST performance, which reflects working memory capacity, declines with age.^57, 68^

**Figure 2.**
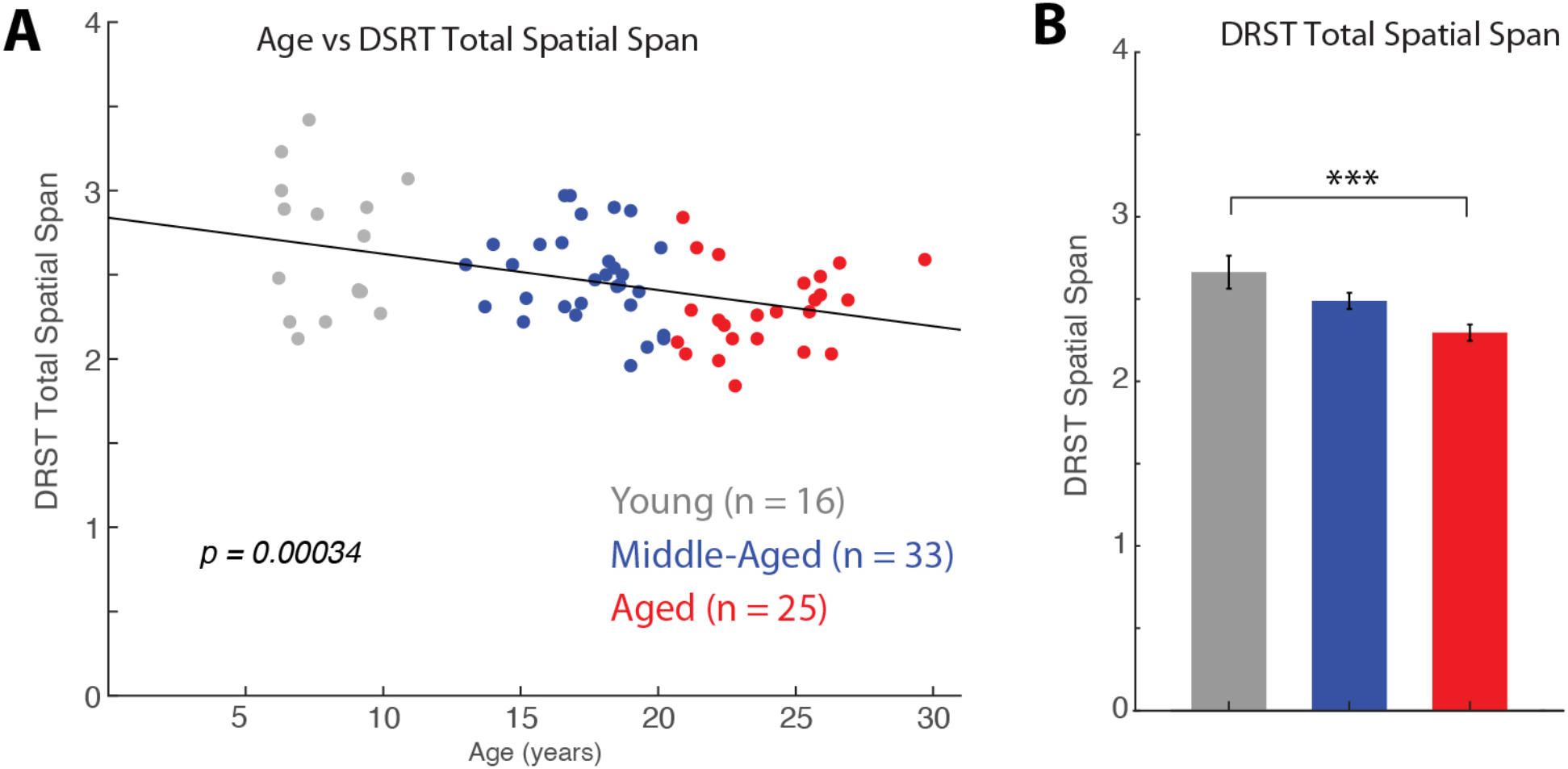
DRST spatial span differences across age. A) Linear relationship between age and DRST spatial span; B) Mean ± SEM DRST spatial span in young, middle-aged and aged monkeys. ***p=0.0008

**Table 2.** Linear Regression Analyses of Age and Measures of Cognitive Task Performance.

| Measurement | Equation | $R^2$ | adjusted $R^2$ | F statistic | p value |
| --- | --- | --- | --- | --- | --- |
| DRST Spatial Span | span = $-0.02 \times \text{age} + 2.84$ | 0.17 | 0.16 | $F(1,69) = 14.3$ | $3.4 \times 10^{-4}$ |
| Discrimination Task - Trials | trials = $-4.40 \times \text{age} + 173.81$ | 0.058 | 0.045 | $F(1,71) = 4.33$ | 0.041 |
| Discrimination Task -Errors | errors = $-2.32 \times \text{age} + 76.83$ | 0.071 | 0.058 | $F(1,71) = 5.38$ | 0.023 |
| CSST Total trials for first category (red) | trials = $9.98 \times \text{age} + 124.63$ | 0.115 | 0.103 | $F(1,73) = 9.34$ | 0.0031 |
| CSST Total errors for first category (red) | errors = $4.25 \times \text{age} + 54.28$ | 0.097 | 0.085 | $F(1,73) = 7.78$ | 0.0068 |
| CSST Total trials for three shift conditions | trials = $39.22 \times \text{age} + 573.26$ | 0.23 | 0.22 | $F(1,73) = 21.3$ | $1.6 \times 10^{-5}$ |
| CSST Total errors for three shift conditions | errors = $19.77 \times \text{age} + 321.93$ | 0.21 | 0.19 | $F(1,73) = 18.7$ | $4.8 \times 10^{-5}$ |
| CSST Total perseverative errors | p-errors = $18.41 \times \text{age} + 201.74$ | 0.29 | 0.28 | $F(1,73) = 29.9$ | $6.3 \times 10^{-7}$ |
| CSST Total perseverative errors as a percent of shift trials | p-errors = $0.23 \times \text{age} + 37.72$ | 0.07 | 0.06 | $F(1,73) = 5.44$ | 0.022 |
| CSST Total number of broken sets | broken sets = $0.30 \times \text{age} + 6.55$ | 0.114 | 0.102 | $F(1,73) = 9.26$ | 0.0033 |
| CSST Total non-responses | non-responses = $2.05 \times \text{age} + 63.62$ | 0.008 | -0.006 | $F(1,73) = 0.56$ | 0.46 |

Based on this analysis and our previous studies, we next grouped the monkeys into categorical age groups (young: 5-12 years, middle-aged: 13-20 years, and aged: >20 years) to better assess when declines in working memory begin and whether there is a significant difference between middle-aged and aged monkeys on the DRST. A one-way ANOVA revealed a significant effect of age group on the total DRST spatial span (Fig. 2B; F(2,67) = 7.61, **p = 0.001)**. Post-hoc tests showed that the total DRST spatial span was significantly greater in young compared to aged monkeys (Tukey’s post hoc test: young vs aged groups, **p = 0.0008**) but the differences between the young and middle-aged monkeys was not significant (**p = 0.145**) although the difference between the middle-aged and aged monkeys approached significance (**p = 0.052**).

Together, these analyses show that there is a gradual age-related decline in performance in working memory, that is greatest in the aged monkeys.

### Discrimination Task Performance

Immediately prior to administering the CSST, monkeys first completed a simple three choice discrimination task for familiarization with the automated testing apparatus and to determine that they could discriminate among multiple stimuli on a screen. Analyses using age as a continuous variable, showed that there was a significant effect of age on both trials and errors to criterion (Table 2, trials: F(1,71) = 4.33, **p = 0.041**; errors: F(1,71) = 5.38, **p = 0.023**). A follow-up one-way ANOVA analysis, with the monkeys grouped by age showed that there was a significant difference between the young and middle-aged monkeys on trials (F(2,69) = 4.44, **p = 0.015**; Tukey’s post hoc test, **p = 0.011**) and errors (F(2,69) = 4.96, **p = 0.0097**; Tukey’s post hoc test, **p = 0.007**) to criterion but no difference between young and aged monkeys or between middle-aged and aged monkeys. While these findings differ slightly from our previous publications that showed no effect of age on this task,^8, 9^ all monkeys reached criterion on this task and therefore we were confident that they were able to discriminate amongst several stimuli and therefore able to complete the CSST.

### Category Set Shifting Task Performance

Similar to the DRST, linear regression analyses with age as a continuous variable, revealed that performance on the CSST declines with age (Fig. 3A-F; See Table 2 for the detailed results). Specifically, we found that the total trials and errors to criterion for abstraction of the first category (red) and the total trials and errors to criterion on the three shift conditions increased with age (Fig. 3A,B; for p values see Table 2). Similarly, the total perseverative errors also showed a significant increase with age (Fig. 3C; **p = 6.3 × 10**^**-7**^) and this effect persisted when expressing the total perseverative errors as a percent of shift trials (Fig. 3D; **p = 0.022**). In addition, the total number of broken sets increased with age (Fig. 3E; **p = 0.0033**). However, there was no significant effect of age on the total number of non-responses (Fig. 3F; **p = 0.46**).

**Figure 3.**
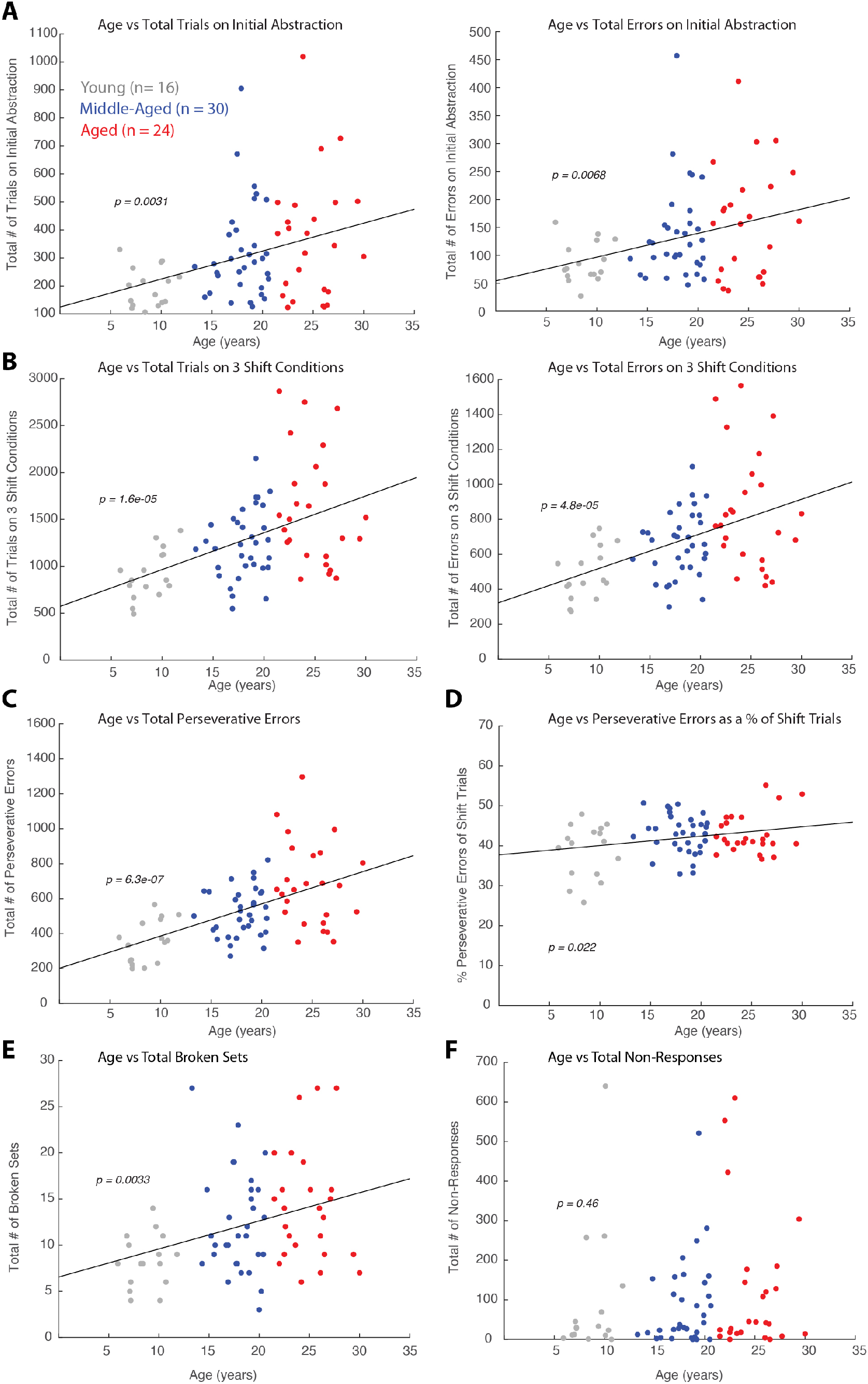
Relationship between CSST performance and age. Linear regression analyses showing relationship: A) Total trials and errors in the initial abstraction (red condition) vs. age; B) Total trials and errors (in three shift conditions) vs age; C) Total perseverative errors vs age; D) Perseverative errors as a percent of shift trials vs age; E) Total broken sets vs age; F) Number of non-responses vs age.

As with the DRST, data was next analyzed with monkeys categorized in three discreet age groups (young: 5-12 years, middle-aged: 13-20 years, and aged: >20 years). Fig. 4A showed that aged monkeys required significantly more trials and made more errors to criterion on the initial abstraction (red) than the young monkeys (Fig. 4A; trials, F(2,71) = 4.64, **p = 0.013**; Tukey’s posthoc test: young vs aged, **p = 0.01**; errors, F(2,71) = 3.45, **p = 0.037**; Tukey’s posthoc test: young vs aged, **p = 0.03**). However, there were no significant differences between the young and middle-aged (trials, **p = 0.058**; errors, p = 0.131) and the middle-aged and aged (trials, p = 0.616; errors, **p = 0.648**) on acquiring the 1^st^ category (red) on the CSST.

**Figure 4.**
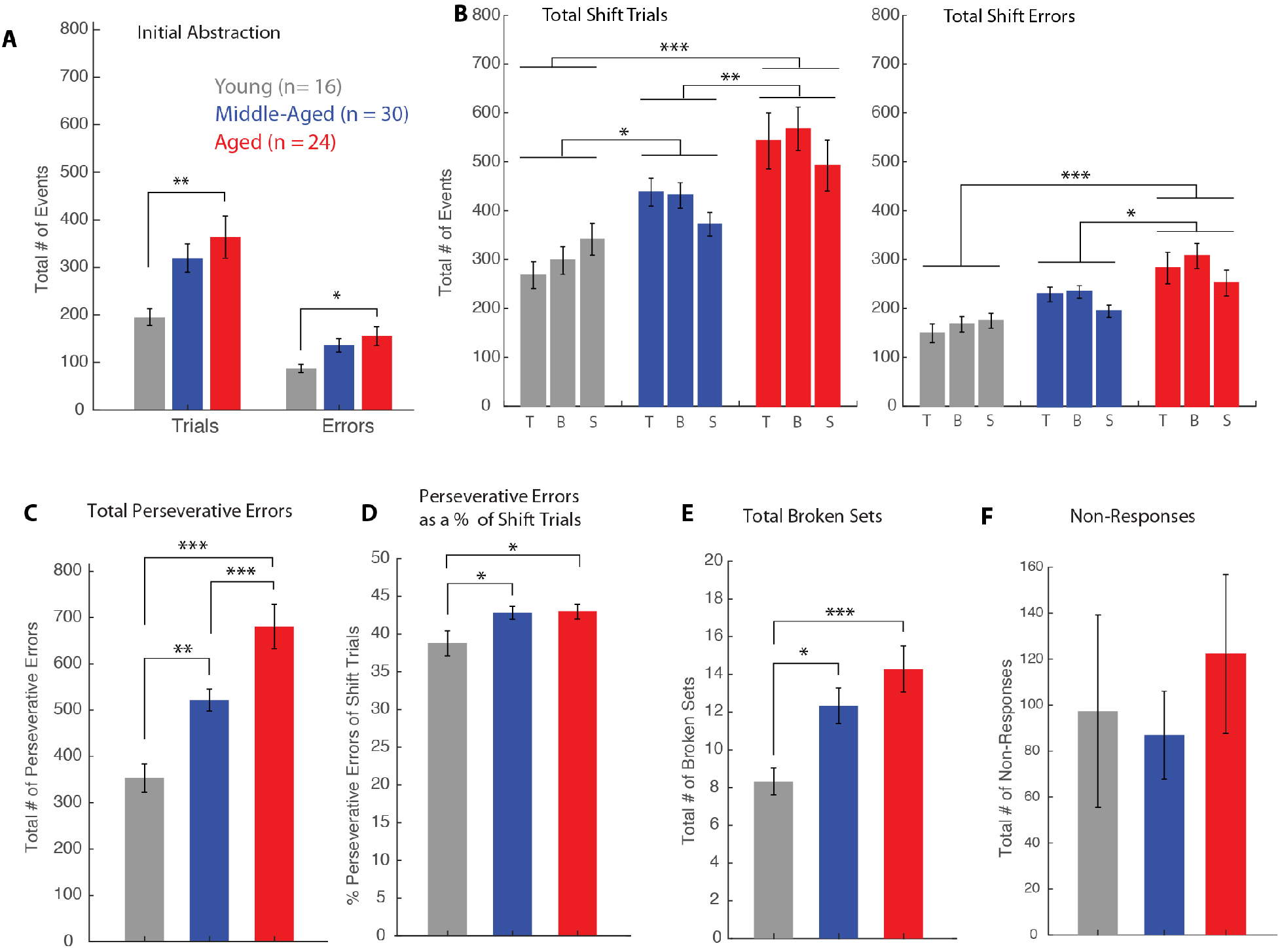
Comparison of CSST performance between young, middle-aged and aged monkeys. Mean ± SEM in young, middle-aged and aged monkeys of: A) Number of trials and errors in the initial abstraction (red condition); B) Number of trials and errors in three shift conditions; C) Number of total perseverative errors; D) Perseverative errors as a percent of shift trials; E) Total Broken sets; F) Number of non-responses. *p < 0.05; **p < 0.01; ***p < 0.001

Examination of the total trials to criterion across the three shift conditions with a two-way ANOVA including age group and condition, with condition as a repeated measure, showed a significant effect of age group (F(2,71) = 12.1, **p = 3 × 10**^**-5**^) and no significant effect of condition (**p = 0.45**) and group by condition interaction (**p = 0.16**). Tukey’s posthoc test revealed a significant difference between the young and middle-aged groups (**p =0.044**), the young and aged groups (**p =2.12 × 10**^**-5**^), and the middle-aged and aged groups (**p = 0.009**) (Fig. 4B).

Similarly, for total errors to criterion across the three shift conditions, there was a significant effect of age group (F(2,71) = 10.7, p = 8.8 **x 10**^**-5**^), no significant effect of condition (**p = 0.087**) and no significant group by condition interaction effect (**p = 0.29**). Tukey’s posthoc test revealed a significant difference between the young and aged groups (**p =6.56 × 10**^**-5**^), and the middle-aged and aged groups (**p = 0.014**) (Fig. 4C). Interestingly, the difference in total errors between young and middle-aged monkeys was the smallest on the final category star (Fig. 4B), which suggests that by the third shift, middle-aged monkeys were learning the shift rule, an ability not seen in the oldest monkeys.

For total perseverative errors, a one-way ANOVA showed a significant effect of age group (Fig. 4D; F(2,71) = 17.07, **p = 8.8 × 10**^**-7**^). A follow-up with the Tukey’s post hoc test revealed a significant difference between the young and middle-aged groups (**p = 0.007**), the young vs aged groups (**p = 5.2 × 10**^**-7**^) and the middle-aged vs aged groups (**p = 0.003**).

Since middle-aged and aged monkeys required more trials to reach criterion, they had more opportunity to perseverate. Therefore, we looked at total perseverative errors a percent of sift trials. Examination of total perseverative errors as a percent of shift trials revealed a significant difference between both young and middle-aged, and young and aged monkeys, but not between the middle-aged and aged monkeys (Fig. 4E; F(2,71) = 3.77, **p = 0.028**; Tukey’s post-hoc test: young vs aged, **p =0.04**; middle-aged vs aged, **p = 0.04**).

Analysis of the total broken sets also showed a significant difference between young and middle-aged monkeys and young and aged monkeys, but not between the middle-aged and aged monkeys (Fig. 4F; F(2,71) = 6.37, **p = 0.003**; Tukey’s posthoc test: young vs. middle-aged; **p = 0.037**; young vs aged **p = 0.002**). Finally, analysis of total non-responses showed no significant differences across age groups (Fig. 4G; F(2,71) = 0.42, **p = 0.66**).

Overall, these findings show age-related impairments in abstraction, set shifting and an increased tendency to perseverate and this is consistent with prior findings with a smaller group of monkeys^8, 9^ and with findings from human studies.^21, 69-71^ Further, there is evidence that significant impairments in perseveration and maintaining a response pattern begins in middle-age and continues to decline with advanced age in the rhesus monkey.

### Age-Related Changes in Passive and Active Physiological Properties of LPFC Pyramidal Neurons

The relationship between age, performance on the DRST and CSST and neuronal properties was analyzed using data from a subset of behaviorally tested monkeys which also had electrophysiological data (n = 19, 18 with both DRST and CSST and 1 with just CSST, Table 1). Some cases from this cohort were also part of the datasets used in previous studies.^39, 40, 57^ Figure 5A depicts the experimental design and workflow of harvesting a fresh tissue block from LPFC area 46, within the ventral bank of the caudal principal sulcus, of cognitively-characterized monkeys. *In vitro* whole cell patch clamp experiments were conducted in 300 μm slices from these fresh tissue biopsies, in order to assess the biophysical properties of layer 3 (L3) pyramidal neurons.

**Figure 5.**
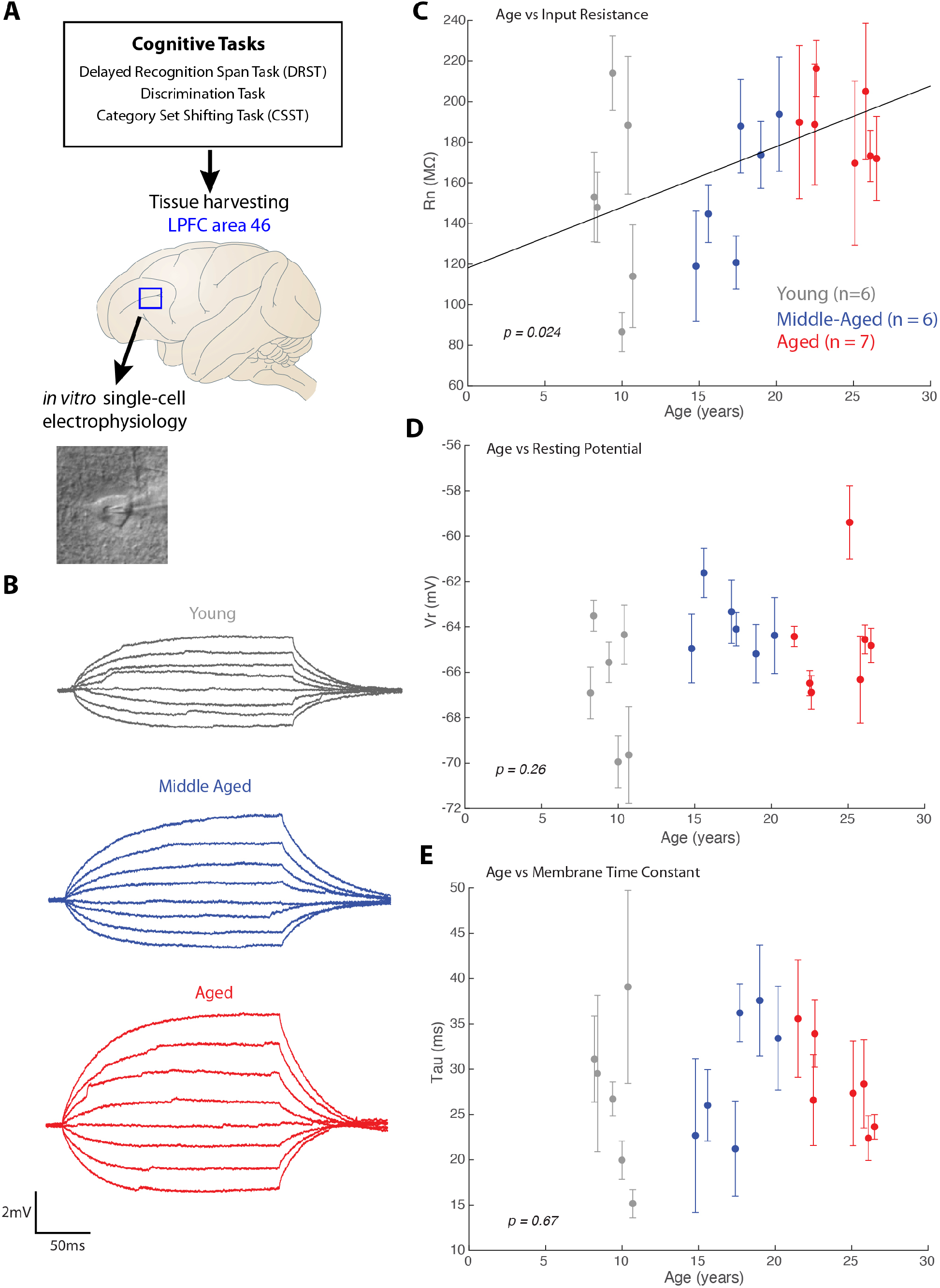
Relationship of age and intrinsic membrane properties of LPFC layer 3 pyramidal neurons. A) Schematic of experimental design of tissue harvesting from LPFC of cognitively characterized monkeys to assess properties of layer 3 pyramidal neurons. B) example voltage traces from which intrinsic properties, Rn and tau were measured; C-E) Linear regression and scatter plot (with error bars indicating ± SEM of cells from each subject) of L3 pyramidal neuron intrinsic membrane properties and age: C) age plotted against mean input resistance (Rn) of neurons from each subject showed a significant positive linear correlation. D) age plotted against mean resting potential (Vr) from each subject and E) age plotted against mean membrane time constant (tau) from each subject showed no significant relationships.

Analyses of the cognitive data from this subset of 19 monkeys was consistent with our analyses from the larger dataset (Fig. 2-4). In particular, the total DRST spatial span for this smaller subset of monkeys showed a significant negative correlation with age (n =18, linear regression, DRST span = -0.02*age + 2.77, *R*^2^= 0.28, adjusted *R*^2^= 0.24, F(1,17) = 6.26, **p = 0.024**). Similarly, there was a significant positive correlation with age for the total preservative errors in this subset of monkeys (n =19; linear regression, perseverative errors = 15.92*age + 260.13, *R*^2^= 0.26, adjusted *R*^2^= 0.22, F(1,18) = 6.08, **p = 0.025**). While the relationship of age with errors on the initial abstraction (red) condition (*R*^2^ = 0.15, adjusted *R*^2^ = 0.10, F(1,18) = 3.04, **p = 0.099**), and total errors on the 3 shift conditions (*R*^2^ = 0.18, adjusted *R*^2^ = 0.13, F(1,18) = 3.66, **p = 0.073**) did not reach statistical significance, these regressions approached significance, consistent with the larger behavioral dataset (Fig. 3). Also consistent with the larger dataset, there was no significant correlation between age and the number of non-responses (F(1,18) = 0.48, **p = 0.496**).

Consistent with previous electrophysiology and aging studies in LPFC,^39, 40, 57^ this study revealed age-related changes in LPFC pyramidal neuron key biophysical intrinsic and firing properties that contribute to increased excitability with age. Specifically, using linear mixed-effects models with random effects and age as a continuous variable, there was a positive linear relationship between age and membrane input resistance (Rn). Higher Rn (greater excitability) was associated with an increase in age (fixed effects regression, Rn = 2.99*age + 118.09, *R*^2^ = 0.13, adjusted *R*^2^ = 0.13, **p = 0.024**; Fig. 5C). However, as we have shown previously^39, 40, 57^, no significant relationship was found between age and resting potential (Vr, **p = 0.26**; Fig. 5D) or membrane time constant (tau, **p = 0.67**; Fig. 5E).

Consistent with the age-related changes observed in input resistance, examination of action potential (AP) firing properties revealed increased excitability of LPFC L3 pyramidal neurons with age. In particular, we found a significant age-related decrease in rheobase current, the minimum current to elicit an AP (fixed effects regression, rheobase = -2.9*age + 140.04, *R*^2^ = 0.23, adjusted *R*^2^ = 0.22, **p = 0.015**; Fig. 6A, B). Thus, with increasing age less current is needed to elicit an AP in LPFC L3 pyramidal neurons. We then examined the repetitive AP firing properties of LPFC neurons in response to 2-sec current injections (Fig. 6C-F). Consistent with our previous findings^39, 40, 57^, for this cohort of monkeys, increasing age was significantly correlated with higher AP firing rates in response to 2-sec current injections (Fig. 6D: I = +130 pA, fixed effects regression, firing rate = 0.38*age + 4.75, *R*^2^ = 0.37, adjusted *R*^2^ = 0.37, **p = 0.004**; not shown: +180pA, fixed effects regression, firing rate = 0.34*age + 8.54, *R*^2^ = 0.36, adjusted *R*^2^ = 0.36, **p = 0.028**. For other currents the effect was not significant).

**Figure 6.**
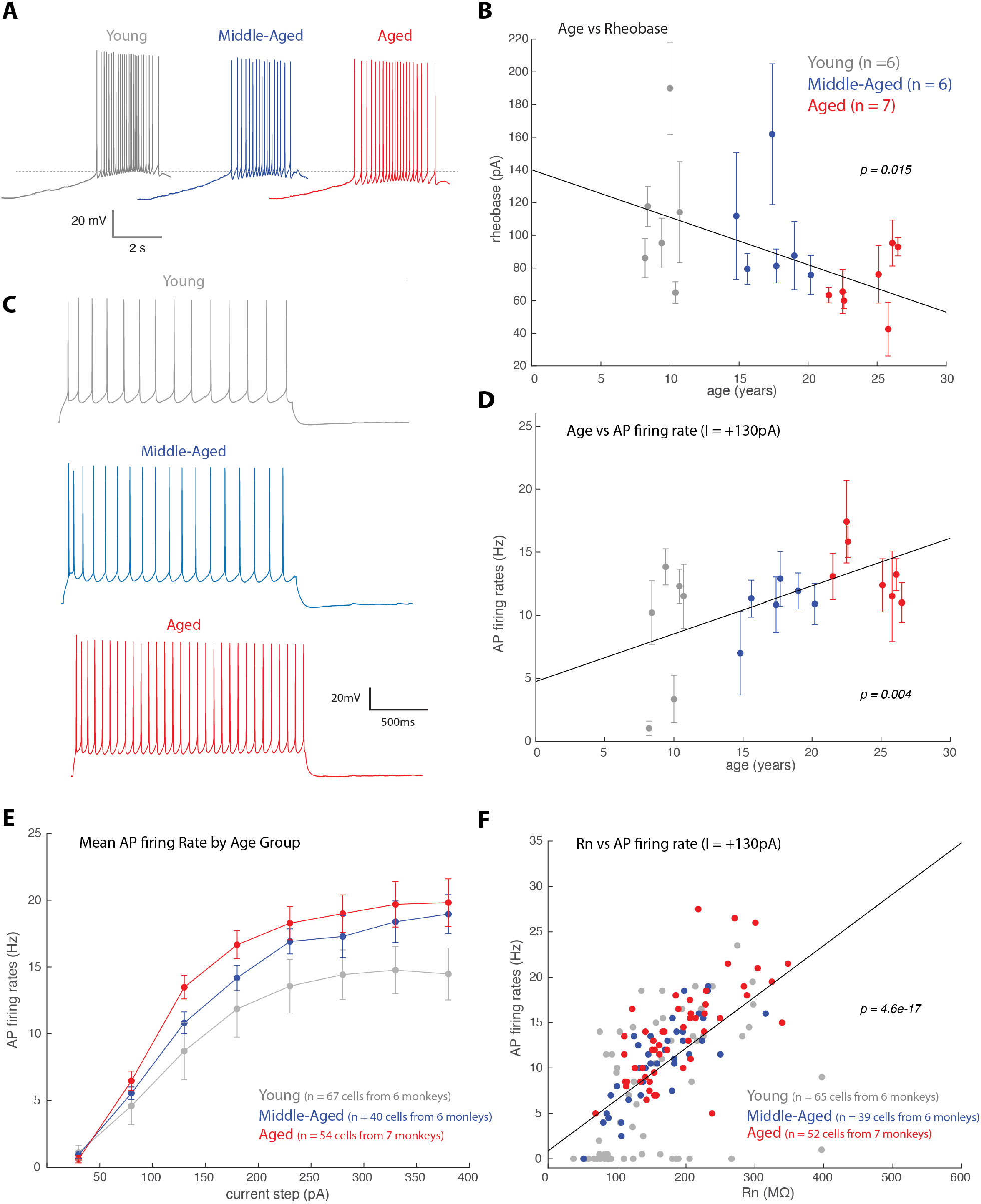
Relationship of age and firing properties of LPFC layer 3 pyramidal neurons. A) Examples of voltage trace showing a train of action potential in response to a current ramp stimulus, which was used to measure rheobase -the minimum amount of current to elicit an AP. The dotted line represents the voltage at which an AP was generated at rheobase in the young monkey group; B) Linear regression and scatter plot (with error bars indicating ± SEM) of age against mean Rheobase of L3 LPFC pyramidal neurons from each subject, showing a significant negative correlation (fixed effects regression, rheobase = -2.9*age + 140.04, *R*^2^ = 0.23, adjusted *R*^2^ = 0.22, **p = 0.015**). C) Example voltage traces showing an AP train in response to a +130pA 2s current step. D) A plot showing a significant correlation between age and AP firing frequency in response to +130pA current injection (fixed effects regression, firing rate = 0.38*age + 4.75, *R*^2^ = 0.37, adjusted *R*^2^ = 0.37, **p = 0.004**). An age-related increase in AP firing frequency was found, which was consistently found in other current amplitudes (not shown). E) Plot of the mean AP frequency in response to a given current amplitude for young, middle-aged and aged monkeys. F) Significant linear correlation between Rn and AP firing frequency in response to +130pA 2s current pulse (linear regression, firing rate = 0.06*Rn + 0.86, *R*^2^ =0.38, adjusted *R*^2^ = 0.37, F(1,151) = 90.38, **p = 4.6 × 10**^**-17**^).

Next, biophysical properties across the three discrete age groups (young, middle-aged and aged) were compared. We found that compared to young monkeys, both middle-aged and aged monkeys had significantly greater mean AP firing rates in response to 2-sec current injections (Fig. 6E). A two-way repeated measures ANOVA, including age group and current step, with injected current as the repeated measure, revealed a significant main effect of age group (F(2,13) = 4.08, **p = 0.042**), injected current (F(7,91) = 175.68, **p = 5.3 × 10**^**-50**^) and interaction of injected current by age group (F(14,91) = 2.25, **p = 0.01**). Moreover, Tukey’s post-hoc test revealed a significant difference between young and aged groups (**p = 0.035**).

The age-related increase in Rn and AP firing rates is related, as linear regression analyses revealed that AP firing responses to a large range of low to high amplitude currents (+30 to +280pA) was significantly correlated with Rn (Fig. 6F, only shown I = +130pA, linear regression, firing rate = 0.06*Rn + 0.86, *R*^2^ =0.38, adjusted *R*^2^ = 0.37, F(1,151) = 90.38, **p = 4.6 × 10**^**-17**^; for all currents from +30 to +230pA, p < 0.001; for +280pA, **p = 0.044**). This confirms the dependence of AP firing on passive membrane properties^64^.

### LPFC Neuronal Firing Rates and Cognitive Performance

Our previous work has shown significant correlations between LPFC neuronal properties and DRST task performance.^57^ Here we build on these findings to assess whether LPFC neuron biophysical properties are related to DRSTsp performance in this group of monkeys and whether they similarly are associated with task performance on the CSST. Thus, we employed linear regression and correlation analyses of electrophysiological and cognitive measures. Consistent with our previous findings,^40, 57^ we found a significant negative linear correlation between DRST spatial span and AP rates in response to +80pA to +280pA current amplitudes (Fig. 7A, I = +80pA, *R*^2^ = 0.37, **p = 0.0079**; +130pA, *R*^2^ = 0.57, **p = 0.0003**; +180pA, *R*^2^ = 0.57, **p = 0.00031**; +230pA, *R*^2^ = 0.47, **p = 0.0018**; +280pA, *R*^2^ = 0.27, **p = 0.027**; see Table 3 for detailed results).

**Figure 7.**
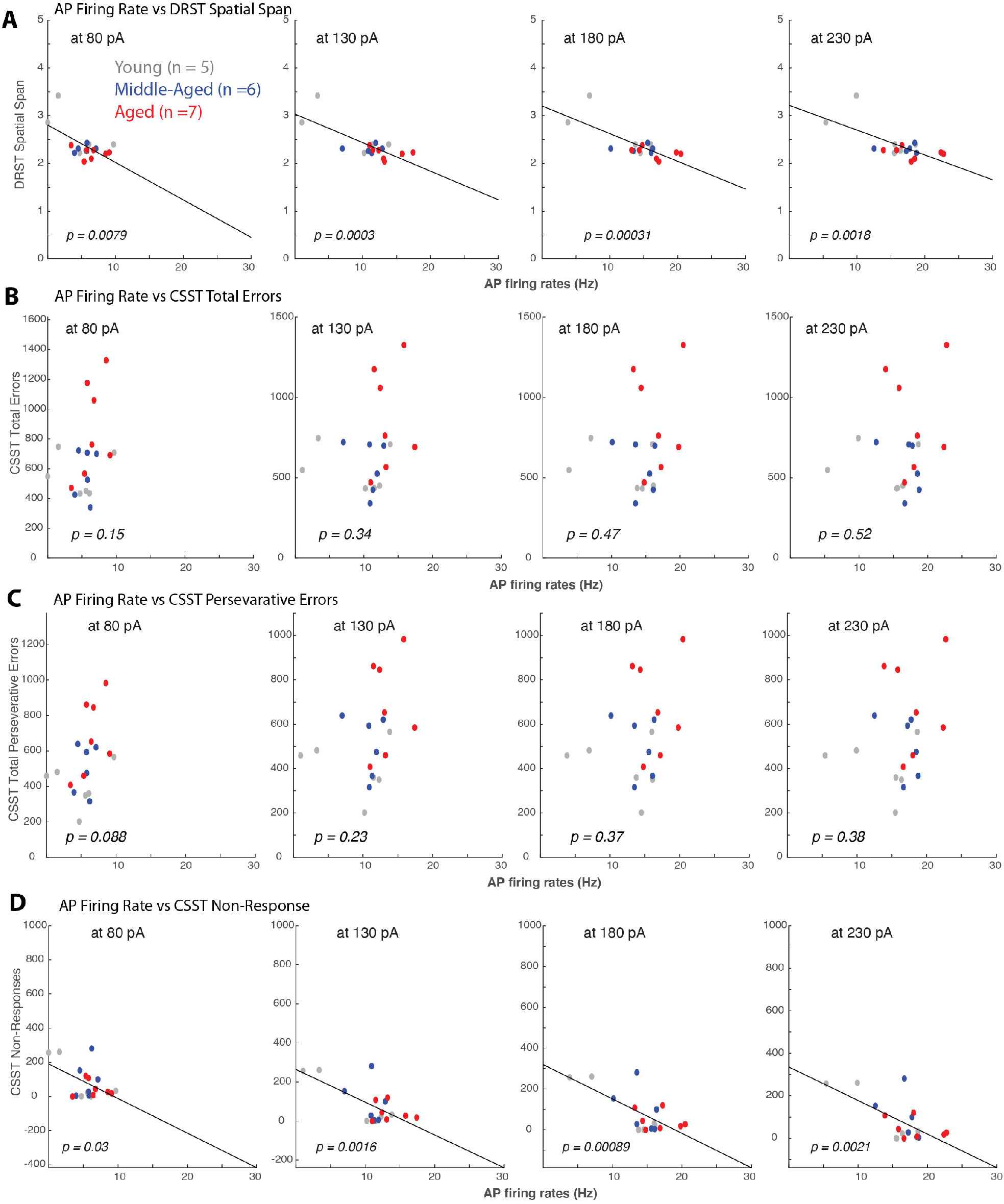
Action Potential Firing rates strongly correlated with DRST but not CSST variables. Linear regression and scatter plots of behavioral variables and mean AP firing rates in response to +80pA, +130pA, +180pA and +230pA current injections of L3 LPFC pyramidal neurons from each subject. A) Significant linear correlations between DRST spatial span and AP rates at all current stimuli amplitudes (I = +80pA, *R*^2^ = 0.37, **p = 0.0079**; +130pA, *R*^2^ = 0.57, **p = 0.0003**; +180pA, *R*^2^ = 0.57, **p = 0.00031**; +230pA, *R*^2^ = 0.47, **p = 0.0018**; +280pA, *R*^2^ = 0.27, **p = 0.027**). Lower DRST spatial span (worse performance) was associated with higher AP firing rates. B-C) Linear regression analyses showed no significant relationships (p > 0.05) between CSST variables: total errors (B) and total perseverative errors (C) and AP firing rates. D) The only CSST variable that showed a significant relationship with LPFC neuron properties were CSST non-responses (I= +80pA, *R*^2^ = 0.25, **p = 0.03**; +130pA, *R*^2^ = 0.45, **p = 0.0016**; +180pA, *R*^2^ = 0.49, **p = 0.00089**; +230pA *R*^2^ = 0.44, **p = 0.0021**). A decrease frequency of non-responses was associated with higher AP firing rates.

**Table 3.** Linear Regression Analyses of LPFC neuron AP firing rate and Measures of Cognitive Task Performance.

| Measurement | Equation | $R^2$ | adjusted $R^2$ | F statistic | p value |
| --- | --- | --- | --- | --- | --- |
| DRST Spatial Span vs Firing Rate (FR) |  |  |  |  |  |
| +80pA | span = $-0.08 \cdot \text{FR} + 2.80$ | 0.37 | 0.33 | $F(1,17) = 9.22$ | 0.0079 |
| +130pA | span = $-0.06 \cdot \text{FR} + 3.03$ | 0.57 | 0.54 | $F(1,17) = 21.12$ | 0.0003 |
| +180pA | span = $-0.06 \cdot \text{FR} + 3.20$ | 0.57 | 0.54 | $F(1,17) = 20.98$ | 0.00031 |
| +230pA | span = $-0.05 \cdot \text{FR} + 3.22$ | 0.47 | 0.43 | $F(1,17) = 13.94$ | 0.0018 |
| +280pA | span = $-0.037 \cdot \text{FR} + 3$ | 0.27 | 0.23 | $F(1,17) = 5.97$ | 0.027 |
| CSST Blue Trials vs Firing Rate (FR) |  |  |  |  |  |
| +80pA | trials = $50.8 \cdot \text{FR} + 194.43$ | 0.23 | 0.18 | $F(1,18) = 5.03$ | 0.039 |
| +130pA | trials = $30.03 \cdot \text{FR} + 144.25$ | 0.21 | 0.17 | $F(1,18) = 4.58$ | 0.047 |
| +180pA | trials = $27.19 \cdot \text{FR} + 88.02$ | 0.18 | 0.14 | $F(1,18) = 3.86$ | 0.066 |
| +230pA | trials = $26.93 \cdot \text{FR} + 38$ | 0.18 | 0.14 | $F(1,18) = 3.86$ | 0.066 |
| +280pA, | trials = $21.18 \cdot \text{FR} + 118.26$ | 0.13 | 0.08 | $F(1,18) = 2.55$ | 0.129 |
| CSST Blue Errors vs Firing Rate (FR) |  |  |  |  |  |
| +80pA | errors = $29.78 \cdot \text{FR} + 99$ | 0.25 | 0.20 | $F(1,18) = 5.62$ | 0.03 |
| +130pA | errors = $17.08 \cdot \text{FR} + 75.48$ | 0.22 | 0.17 | $F(1,18) = 4.72$ | 0.044 |
| +180pA | errors = $15.54 \cdot \text{FR} + 42.44$ | 0.19 | 0.14 | $F(1,18) = 4.03$ | 0.061 |
| +230pA | errors = $15.23 \cdot \text{FR} + 16.49$ | 0.19 | 0.14 | $F(1,18) = 3.92$ | 0.064 |
| +280pA, | errors = $12.12 \cdot \text{FR} + 59.35$ | 0.13 | 0.085 | $F(1,18) = 2.66$ | 0.121 |
| CSST Blue Perseverative Errors vs Firing Rate (FR) |  |  |  |  |  |
| +80pA | p-errors = $23.28 \cdot \text{FR} + 91.53$ | 0.26 | 0.22 | $F(1,18) = 6.11$ | 0.024 |
| +130pA | p-errors = $13.65 \cdot \text{FR} + 69.75$ | 0.24 | 0.197 | $F(1,18) = 5.42$ | 0.032 |
| +180pA | p-errors =<br>12.56*FR + 41.24 | 0.22 | 0.17 | F(1,18) = 4.75 | 0.044 |
| +230pA | p-errors =<br>12.26*FR + 21.22 | 0.21 | 0.16 | F(1,18) = 4.56 | 0.048 |
| +280pA, | p-errors =<br>9.55*FR + 59.32 | 0.15 | 0.096 | F(1,18) = 2.92 | 0.106 |
| CSST Non-Responses vs Firing Rate (FR) |  |  |  |  |  |
| +80pA | non-responses = -<br>20.25*FR +<br>190.75 | 0.25 | 0.20 | F(1,18) = 5.63 | 0.03 |
| +130pA | non-responses = -<br>16.76*FR +<br>264.08 | 0.45 | 0.42 | F(1,18) =<br>14.10 | 0.0016 |
| +180pA | non-responses = -<br>16.83*FR +<br>319.24 | 0.49 | 0.46 | F(1,18) =<br>16.14 | 0.00089 |
| +230pA | non-responses = -<br>15.79*FR +<br>335.86 | 0.44 | 0.40 | F(1,18) =<br>13.19 | 0.0021 |
| +280pA, | non-responses = -<br>13.76*FR +<br>311.62 | 0.39 | 0.34 | F(1,18) =<br>10.35 | 0.005 |

In contrast to DRST, linear regression analyses showed no significant relationships between the measures of CSST performance and any of the LPFC biophysical measures (Fig. 7B, C). For example, AP firing rates did not correlate with the total errors and total perseverative errors on the CSST (Fig. 7B, AP firing vs total errors: **p > 0.05** at all injection levels; Fig. 7C, AP firing vs total perseverative errors: **p > 0.05** at all injection levels). Although there were two exceptions. First, there was a significant correlation of AP firing rates in response to low to medium amplitude input currents with trials, errors, and perseverative errors in the blue shift condition (Table 3). Because the correlation was only found with one shift condition, this finding is difficult to interpret. Second, there was a strong negative correlation between AP firing rates in response to +80pA to +380pA stimulus amplitudes and the total non-responses on the CSST (Fig. 7D; non-responses vs AP firing frequency at: I= +80pA, *R*^2^ = 0.25, **p = 0.03**; +130pA, *R*^2^ = 0.45, **p = 0.0016**; +180pA, *R*^2^ = 0.49, **p = 0.00089**; +230pA, *R*^2^ = 0.44, **p = 0.0021**; Table 3). However, as noted in the previous section, there is no significant relationship of non-responses with age (Fig. 3F and 4F) and again this finding is difficult to interpret.

Our findings are consistent with the hypothesis that aging leads to increases in AP firing rates in PFC which in turn reduces DRST task performance (but not CSST task performance). We used mediation analysis (Methods) to test this hypothesis. For I = +180 pA we found a reduction of the direct effect of age on DRST span when including the indirect effect of age × firing rate (from β = -0.0243, p = 0.0453 to β = -0.0085, p = 0.1827). The indirect effect was significant (β = -0.0158, p = 0.0488), demonstrating that the decrease in DRST span with age was mediated by the change in prefrontal AP firing rate. Similar results were obtained for AP firing rates measured other stimulation currents in the intermediate range though p-values were slightly above 0.05 (I = 130 pA: p = 0.0732, I = 230 pA: p = 0.0547). Finally, consistent with the lack of correlation between AP firing rate and CSST task performance, we observed no mediation effect for the total CSST errors and perseverative errors (I = +180 pA, p = 0.9075 and p = 0.9260, respectively).

## Discussion

### Summary of Results

The current study used a large dataset allowing for direct comparison of the effects of age as a continuous variable on two distinct cognitive domains – working memory assessed with the DRST and executive function assessed with the CSST in the same subjects. Additionally, in a subset of these subjects measurements of biophysiological properties of layer 3 neurons in the LPFC enabled examination of the relationship between performance on these cognitive tasks and biophysical properties of LPFC neurons. These analyses revealed that: **First**, consistent with previous data,^40^ **DRST** spatial memory span showed a significant negative linear relationship with age. Comparisons between categorical age groups showed that aged monkeys but not middle-aged monkeys, demonstrated significantly lower memory spans than young monkeys. **Second**, with age as a continuous variable, the data revealed that **CSST** performance changed with age, with progressive impairments in set shifting and increased perseverative errors showing the strongest correlations with age. Comparisons between age groups further demonstrated that the onset of impairments in EF start in middle-age and become increasingly severe with advanced age. These current findings using a large dataset corroborated our previous findings in a smaller group of monkeys, which showed that relative to young adult monkeys, middle-aged and aged monkeys evidenced significant impairments in abstraction, set-shifting and demonstrated a high degree of perseverative responding.^8, 9^ **Third**, in a subset of monkeys with single-cell electrophysiology data, the analysis revealed age-related changes in L3 LPFC pyramidal neuron properties, namely a significant increase in excitability with age (higher Rn, lower rheobase and increased AP firing frequency). Further, this age-related hyperexcitability of LPFC neurons was significantly correlated with age-related decline in DRSTsp, but not with CSST performance measures.

### Age-Related Changes in Working Memory and Executive Function

The findings in the current study and our previous work demonstrate that aged monkeys show significant impairment on a task of spatial working memory. ^10, 17, 68^ Specifically, performance on the DRSTsp showed that spatial span decreased with age reflecting an age-associated impairment in spatial working memory. However, when categorical comparisons were made, the significant difference was observed only between aged and young monkeys with aged monkeys achieving shorter spans of correct responses than young monkeys while there was no significance difference between the middle-aged and young monkeys. On the other hand, consistent with our previous work, both middle-aged and aged monkeys were impaired on the CSST, a task that required them to abstract a category based on reward contingencies, maintain that response pattern until a change in reward contingency occurred and then shift their response pattern accordingly. ^8, 9^ On the CSST, most of the variables, specifically the total errors and perseverative errors, exhibited stronger correlations with age than did the DRSTsp span. Compared to young monkeys, both middle-aged and aged monkeys required more trials and made more errors when learning the initial abstraction category and during all subsequent shift conditions on the CSST. In addition, an error analysis revealed that both the middle-aged and aged monkeys demonstrated a marked tendency to perseverate with the oldest monkeys demonstrating the greatest tendency to perseverate.^8, 9^

Overall, the pattern of impairments on the DRST and CSST demonstrated by the middle-aged and aged monkeys closely resembles impairments observed clinically with aging humans.^11, 69, 72^ In terms of WM, several studies have demonstrated age-related deficits on working memory on tasks such as the Corsi Block Tapping test^73^ and Digit Span^74^, with some evidence for declines beginning as early as the third decade of life, but becoming most prominent after the sixth decade.^4, 5, 7, 75-77^ Consistent with this, we show that age as a continuous variable was significantly correlated with a decline in DRST span. While in the present study the middle-aged monkeys as a group were not significantly impaired relative to young monkeys, their spans overall were lower than the young monkeys (p=0.145). With regard to EF, human clinical studies have established age-related declines in concept formation, abstraction, verbal and non-verbal cognitive switching and mental flexibility and concrete response patterns beginning in middle-age.^1, 69, 78-83^ Aging also negatively affects the ability to inhibit an established response pattern in favor of producing a novel response, a skill necessary for successful completion of the human Wisconsin Card Sorting Task (WCST)^82^ and CSST shown here in monkeys. Taken together, these data suggest that the age-related impairments in spatial WM are mild in middle age and only become severe with advancing age, whereas declines in EF appear to begin in early middle-age and progress in severity with advancing age.

### Cognitive Decline in Middle-age

Until recently many studies of cognitive aging have focused on aged individuals (60+ years). However, recent studies have shown that age-related changes in cognition can occur as early as the 4^th^ and 5th decades and include declines in WM and EF. ^77-79, 84, 85^ For example, Singh-Manoux et al, 2012 ^80^, demonstrated in the Whitehall II study beginning at approximately age 45, individuals show declines in inductive reasoning and verbal and mathematic reasoning tasks.^80^ Further, Salthouse (2009) provided significant evidence of age-related cognitive decline beginning in the 30s in humans with declines occurring earliest in “fluid” intelligence and multi-tasking skills.^76, 78, 79, 84, 86, 87^ Also, most relevant to the current study, a review of normative data on the WCST in humans shows that not only are individuals of advanced age less efficient on this task with evidence of impaired response maintenance and shifting and perseveration, but these impairments begin to occur in middle age. ^69^

The results in the current study show that middle-aged monkeys were not significantly impaired on the DRST but were significantly impaired on the CSST provides evidence that EF may be among the earliest domains of cognitive function to exhibit change with aging. A striking aspect of the deficit displayed by many of the middle-aged and aged monkeys was the frequency of perseverative errors. This deficit worsens gradually with aging, as indicated by a linear increase in perseverative errors with increasing age. A similar pattern of results was shown in a study examining WSCT in humans, which showed an increase in perseverative errors on the WCST by both middle aged and aged individuals and a strong correlation between age and perseverative errors. ^69^

### Age-Related Impairments in WM and EF are Similar in Nature to Cognitive Performance Deficits Following PFC Lesions

While age related changes in WM and EF are well established in humans and this study demonstrates a similar pattern in a large group of male and female rhesus monkeys, the neurobiological basis of these functions is not well understood. However, there is substantial evidence that both WM and EF are mediated by the PFC. ^88-100^ In a separate study in our laboratory we showed that young monkeys with lesions encompassing several areas in the PFC (areas 46, 8, 9 and 10^93^ and unpublished data) were impaired on both the DRST and CSST. Interestingly, on both tasks, the performance of the young monkeys with PFC lesions on both tasks was similar in nature to the performance of the middle-age and aged monkeys in the current study. Specifically, these animals demonstrated shorter memory spans, perseveration and an inability to shift and to use feedback to modify response patterns^93^. Taken together with our findings of hyperexcitability of aged PFC neurons, this data further support the notion that performance on the DRST and CSST relies, at least in part, on the functional integrity of the PFC. This notion is also supported by studies with human subjects that have demonstrated that humans with circumscribed frontal lobe damage are impaired on working memory tasks and the WCST. ^101-103^ Results from studies with both monkeys and humans showing impairments in WM and EF following damage or injury to the PFC provide evidence that age-related declines in these cognitive domains likely result, at least in part, from PFC dysfunction.^99, 103-106^ However, since lesion studies in monkeys and brain trauma in humans result in damage to multiple areas of the PFC, the contributions of distinct PFC areas remains unclear. Further, these large lesions in the PFC result in pathway disconnection and network changes that are likely related to impairments in specific aspects of the WM and EF and therefore need more investigation.

### Age-Related Changes in Intrinsic LPFC Neuron Properties are Associated with Impairments in Spatial Working Memory in Aging

Age-related neural changes in several structural, molecular and functional properties of neurons localized within area 46 of the LPFC have been associated with cognitive decline.^19, 36^-^39, 41, 42, 44-46, 48, 49, 51, 52, 108^ Specifically, decreases in synapses and synaptic transmission, ^38, 41, 42, 45, 51^ and alterations in the expression of many neuromodulatory receptors,^19, 46^ and decreases in dendritic complexity^37, 109^ have been found in LPFC and are correlated with age-related cognitive decline. Following this work, in the present study we examined the relationship between performance on the DRST and CSST and biophysical properties of neurons within LPFC, specifically within area 46. Consistent with previous work,^39, 40, 57^ we have shown here that age-related changes in single-neuron biophysical properties of pyramidal neurons in L3 of LPFC in particular correlate with impaired performance on the DRSTsp, but not CSST. These data suggest a potential dissociation between specific cognitive functions – WM performance may depend more strongly on L3 neurons in LPFC area 46 than performance on the CSST. It is also worth noting that while the majority of CSST measures reflecting the main aspects of EF did not correlate with LPFC neuronal properties, there was a strong correlation of LPFC neuronal firing frequency with CSST non-responses. This could reflect an overall effect of attentional state, consistent with the well-known role of LPFC area 46 in visuospatial attention. However, since no significant relationships were found between non-responses and age and other CSST performance measures, it is difficult to speculate on the relationship between LPFC neuronal properties and CSST non-responses based on our current dataset.

The finding of a relationship between L3 LPFC pyramidal neuron biophysical properties and performance on the DRSTsp but not on the CSST was unexpected since the cognitive abilities necessary to complete the CSST (abstraction, set-shifting and inhibition of perseveration) are well established as mediated by the PFC. However, the PFC is a heterogenous region that consists of distinct but a highly interconnected networks of lateral, medial and orbital cortical areas.^110^ Each of the PFC regions are important for higher-order cognitive functions, and play specific roles in cognition that are not yet fully understood.^100, 111-114^

### LPFC Neuron Properties do not Correlate with CSST Performance: Implications for Age-Related Changes in PFC Networks

Layer 3 pyramidal neurons in LPFC are thought to be an important neuronal substrate for WM. ^115-119^ Layer 3 pyramidal neurons are involved in cortico-cortical communication^109^ and sustained firing activity and recurrent synaptic connectivity between LPFC L3 pyramidal neurons, specifically in area 46, are thought to be necessary to maintain information in working memory.^117, 120-122^ Previous empirical and computational work from our group has shown that the combination of age-related hyperexcitability and decreases in excitatory synaptic strength and activity in L3 LPFC neurons are the strongest predictors of impairments in maintaining spatial information in the DRSTsp.^57^ The specific role of LPFC area 46 (along with area 8, the frontal eye fields) in spatial working memory have also been shown via *in vivo* recordings of task related neuronal activity in monkeys during delayed oculomotor response tasks.^123, 124^ Our data is consistent with previous studies in monkeys and humans that suggest neuronal activity underlying spatial working memory, such as that required for execution of the DRSTsp, is strongly localized to the LPFC.^113, 124, 125^

In contrast, EF tasks that require higher cognitive demand, such as the CSST, may depend more strongly on a larger network of PFC regions. Indeed, previous *in vivo* physiological studies from awake monkeys have suggested that the engagement of rostral, medial and orbital PFC areas as cognitive task demands increase.^123^ The rostral medial PFC and anterior cingulate cortex (ACC), for instance, are thought to be specifically engaged with the LPFC when there are increased attentional demands, conflict and errors, task-switching paradigms^113, 124-133^ and during the feedback and performance evaluation phase of cognitive tasks.^134, 135^ Further, evidence from imaging studies has shown that the fMRI bold activation of the dorsal ACC was the first region to show activity during performance on a Stroop task,^136-138^ a task that assesses aspects of EF such as abstraction, cognitive switching and mental flexibility, similar to the CSST.

Anatomical studies have revealed robust connections between the ACC and the LPFC.^139, 140^ This is interesting especially since age-related myelin damage occurs within the anterior cingulum bundle and the frontal white matter.^23, 34, 53, 55^ Thus, the connectivity between ACC and other PFC regions, including the LPFC, are likely to be the most impacted with age. In addition to age-related frontal and anterior cingulate white matter degeneration seen with age, aging is associated with a marked the loss of synapses and synaptic structures in LPFC area 46. ^36, 38, 40-44^ Our group and others have reported a dramatic loss of spines (∼40% decrease) on L3 LPFC pyramidal neurons associated with age.^36, 38, 39, 41, 42, 44, 141, 142^ Ultrastructural studies have demonstrated a marked age-related decrease in the density of excitatory synapses in L1 and L2-3 of LPFC area 46. ^34, 45, 143^ Further, glutamatergic axonal boutons are structurally altered with age.^108^ Collectively, these data point to significant age-related changes in connectivity within the PFC networks involved in cognition that likely underlie the impaired performance on the DRSTsp and CSST shown in this study.

## Conclusions

This study demonstrates that performance on both the DRST and CSST tasks declines with age and the most pronounced age-related changes occurring in the ability to shift cognitive set and inhibit perseveration, impairments that begin in middle-age. In addition, we demonstrated that performance on the DRST but not CSST was associated with changes in the biophysical properties of LPFC neurons. Our previous studies with localized lesions in PFC encompassing areas 46, 9, 10 and 8 showed impairments in both CSST and DRST(^93^ and unpublished data). Taken together with previous work, our current findings point to the importance of intrinsic LPFC neuronal properties for spatial working memory functions such as required by the DRST. In contrast, for performance on more complex EF tasks such as CSST, deficits only emerge when broader PFC networks are disrupted. Age-related changes in excitability and biophysical signaling properties of LPFC neurons will affect signal propagation via white matter tracts and synaptic transmission to downstream targets. Together with the age-related synapse loss within LPFC and white matter degeneration in frontal white matter tracts,^34, 36, 45, 143^ hyperexcitability of LPFC neurons predicts impaired PFC network signaling dysfunction with age.^57^ Future work will need to focus on the temporal progression of age-related changes within distinct areas and networks in order to dissociate the neural mechanisms underlying specific functional impairments in cognitive aging.

## Acknowledgments

The authors thank Bethany Bowley, Penny Shultz, Karen Slater and Ethan Gaston for their invaluable assistance with this project and the CERCA Program/Generalitat de Catalunya for institutional support. This project was funded by: NIH/NIA RF1AG043640, NIH/NIA RF1AG062831, NIH/NIA R01AG078460, NIH/NIA R01AG043478, NIH/NIA P01 AG000001, NIA-NSF CRCNS R01 AG071230, NIH/NIA R01 AG059028, PCI2020-112035 (Spanish State Research Agency (AEI)), the European Union “NextGenerationEU / PRTR”, the Spanish State Research Agency, Severo Ochoa and Maria de Maeztu program for Centers and Units of Excellence in R&D (CEX2020-001084-M).

